# Evaluating cell-free RNA extraction platforms for RNA liquid biopsy via nanopore sequencing

**DOI:** 10.64898/2026.11.19.767748

**Authors:** Connor S. Mattingly, Alexander D. Hill, Daniel H. Kim

## Abstract

RNA liquid biopsies provide systemic insights into health and disease by profiling cell-free RNAs (cfRNAs) secreted into the circulation. While cfRNAs are thought to be short or fragmented, recent efforts using long-read nanopore sequencing have found that thousands of longer, full-length cfRNAs are highly abundant in plasma. However, cfRNA extraction methods have not been rigorously evaluated to assess reproducibility, cfRNA length, and transcriptome complexity. Here we show that extraction workflow is a major source of variation by comparing identical plasma samples across three different cfRNA extraction platforms. While these platforms recover comparable quantities of cfRNA, we perform nanopore sequencing to show that they differ in their cfRNA composition based on read-length distribution and biotype representation. Principal component analysis and hierarchical clustering of variable protein-coding and repeat-derived cfRNAs show that samples tend to cluster based on extraction platform. Robust detection of specific biotypes is contingent on extraction platform, as evidenced by significant enrichment of Alu element- and long noncoding RNA (lncRNA)-derived cfRNAs in some but not all methodologies. Our results reveal that cfRNA extraction platforms influence the composition and complexity of downstream libraries, highlighting the importance of selecting the right platform for profiling disease-relevant biomarkers in RNA liquid biopsy studies.

## INTRODUCTION

Cell-free RNA (cfRNA) circulating in human blood plasma provides systemic biological information that can be leveraged for disease detection, monitoring, and assessment of treatment response [1–7]. Unlike cell-free DNA such as circulating tumor DNA, which is released largely upon cell death [8,9], cfRNA is actively secreted by living cells and reflects the transcriptional state of its cell of origin [4,10–13]. These transcripts persist in the circulation by being packaged into extracellular vesicles (EVs) [14–16] or through their association with apoptotic bodies and ribonucleoprotein complexes, which protect them from nucleases in the blood [17–19]. cfRNA profiling has relied predominantly on short-read sequencing, making it more challenging to assign reads to unique genomic loci, to identify cfRNA isoforms, and to determine true cfRNA transcript length [20–23]. Long-read sequencing platforms, such as nanopore sequencing, help address these limitations by sequencing full-length cfRNA transcripts and isoforms [24–28].

RNA liquid biopsy platforms benefit from methods that capture and profile full-length cfRNA molecules [29]. However, upstream cfRNA extraction is technically challenging because cfRNA is present at relatively low concentrations in plasma and spans a broad range of transcript lengths [19,30]. cfRNA extraction platforms also use distinct capture methods, such as total nucleic acid-binding magnetic beads or column-based EV capture prior to RNA extraction, which may result in the recovery of varying cfRNA populations [19,31]. Different capture methods have been shown in short-read sequencing studies to alter the composition of recovered cfRNA, but such inherent biases have not been explored using long-read cfRNA profiling approaches [19,31,32]. Only a handful of studies have profiled plasma cfRNA using long-read sequencing [5,27], and cfRNA extraction platforms have not been systematically evaluated to determine their effects on long cfRNA molecule recovery, representation of the cfRNA transcriptome, or technical reproducibility.

Here we benchmark three cfRNA extraction platforms using independent pools of human plasma to assess the biological and technical reproducibility of each workflow using long-read nanopore sequencing. We evaluate several key metrics, including total cfRNA recovery, cfRNA transcript length, and biotype composition and bias. Our study reveals how cfRNA extraction platforms affect the reproducibility and representation of the cfRNA transcriptome for RNA liquid biopsy workflows.

## RESULTS

### Experimental design for evaluating cfRNA extraction platforms

Three independent plasma pools (A, B, and C) served as biological inputs for a matched comparison of three cfRNA extraction platforms (Fig. 1a). Each pool was filtered through a 0.8- µm membrane to remove any platelets or cell debris and divided into input volume-matched aliquots for triplicate cfRNA extractions using the nRichDX Revolution cfTNA Max 20 Kit (nRichDX), the Norgen Plasma/Serum Circulating and Exosomal RNA Purification 96-Well Kit Slurry Format (Norgen), or the Qiagen ExoRNeasy Maxi Kit (Qiagen), yielding nine cfRNA samples in total. Equal aliquots from each of the three independent pools were processed on all three cfRNA extraction platforms, so that each pairwise platform comparison is on matched biological input material. The extraction platforms differed primarily in their cfRNA capture methodology: nucleic acid-binding magnetic beads (nRichDX, Norgen) or column-based EV capture prior to RNA extraction (Qiagen). All cfRNA eluates underwent post-extraction DNase treatment, and cfRNA libraries were then prepared using the same long-read cDNA workflow for subsequent nanopore sequencing.

**Figure 1.**
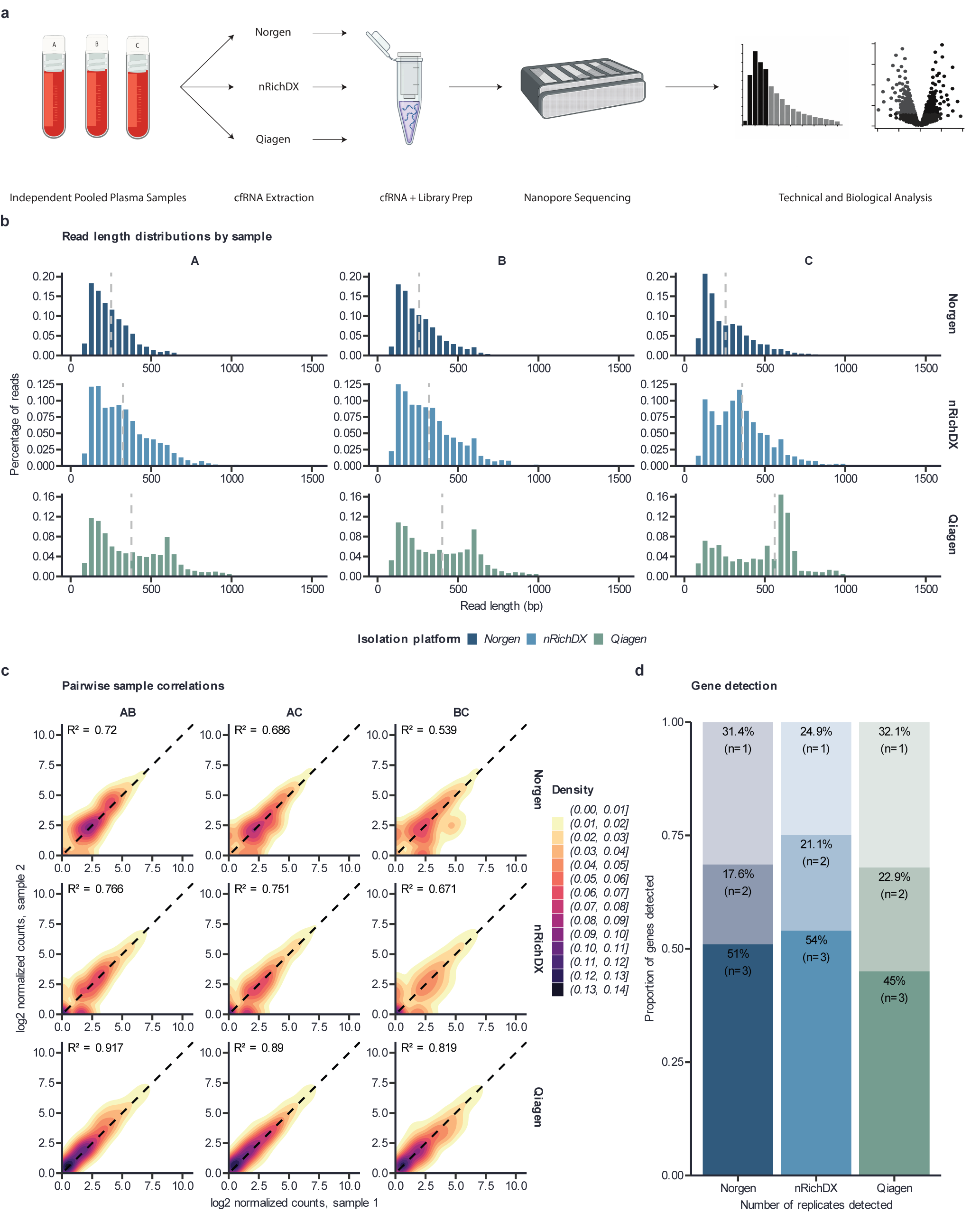
Technical performance of cfRNA extraction platforms. **(a)** Experimental workflow. Three independently pooled plasma samples (a-c) were split into three aliquots. Each aliquot was extracted with a different cfRNA kit: Norgen’s Plasma/Serum Circulating and Exosomal RNA purification 96-Well Kit Slurry Format (Norgen), nRichDX’s Revolution cfTNA Max 20 kit (nRichDX), and Qiagen’s ExoRNeasy Maxi Kit (Qiagen), and sequenced using Oxford Nanopore Technologies’ PromethION platform (9 libraries total). **(b)** Read length distributions by extraction platform (rows) and plasma pool (columns). Bars show the fraction of reads in 50bp bins, and dashed lines mark the N50 read length. **(c)** Pairwise comparison of each plasma pool sample between extraction platforms. Each axis represents each sample’s quantification, and density highlights the aggregate concentration of gene-level quantifications. The dashed line represents perfect correlation, and R^2^ is shown for each pairwise comparison. **(d)** Proportion of genes detected in one, two, or three plasma pools for each platform. A gene is considered at >= 3 counts.

### Technical performance of cfRNA extraction platforms

All three extraction platforms exhibited comparable cfRNA recovery, and from 1.3 mL of input plasma, the mean recovered cfRNA mass was 2.92 nanograms (ng) for Norgen, 2.90 ng for Qiagen, and 2.44 ng for nRichDX. We generated long-read cDNA libraries from each cfRNA sample and performed nanopore sequencing on the PromethION platform. To comprehensively profile the cfRNA transcriptome, including repeat element-derived cfRNAs [5,10,11,33,34], nanopore reads were then aligned to the human genome (GRCh38) using a repeat-aware approach that incorporates the RepeatMasker annotation.

We first characterized cfRNA insert sizes to determine the read length distributions of cfRNA molecules recovered by each extraction platform (Fig. 1b). Across all of the workflows, 66.2 - 80.8% of cfRNA reads were longer than 200 nucleotides (nt), and the 95th percentile ranged from 626 - 960 nt across the platforms. The proportion of cfRNA reads longer than 200 nt was 80% for Qiagen, 77.5% for nRichDX, and 66.2% for Norgen, while the read N50 values were 626 nt for Qiagen, 550 nt for nRichDX, and 396 nt for Norgen.

We next looked for potential internal priming events in our sequencing data to determine where there might be any cell-free DNA contamination, using previously described criteria [35], where internal priming is assumed to potentially occur if the 20 nt sequence downstream of a read is found to contain six consecutive adenosines, or at least seven adenosines within a 10-nucleotide window. Based on these criteria, potential internal priming rates for our cfRNA libraries were 4.20% (± 0.16%) for nRichDX, followed by 4.07% (± 0.44%) for Norgen, and 2.19% (± 0.42%) for Qiagen.

### Reproducibility of transcriptomic representation across cfRNA extraction platforms

We next assessed how reproducibly each extraction platform recovered the cfRNA transcriptome across the three independent plasma pools, comparing per-gene abundance estimates and the identities of the detected gene sets. Mean pairwise replicate concordance (R^2^) was 0.875 for Qiagen, 0.73 for nRichDX, and 0.65 for Norgen (Fig. 1c).

Given that the per-gene count distributions for nRichDX and Norgen were centered higher than for Qiagen (Fig. 1c), we calculated transcripts per million (TPM) for each transcript to compare relative abundances. nRichDX libraries had a mean expression of 681.6 TPM (range 566.6– 768.6 TPM), while Qiagen had a mean of 405.5 TPM (range 366.6–441.3 TPM) and Norgen had a mean of 389.5 TPM (range 344.8–448.2 TPM). Moreover, the extraction platforms also differed in their reproducibility of gene detection across all independent plasma samples, with nRichDx detecting 54% of genes, Norgen detecting 51% of genes, and Qiagen detecting 45% of genes across all independent plasma samples (Fig.1d).

### Extraction platform-dependent cfRNA transcriptome variability

Principal component analysis of variance-stabilized counts revealed that the first principal component (PC) explained 38.88% of the variance, while PC2 explained 18.11% of the variance (Fig. 2a). Independent plasma samples A, B, and C appeared to cluster more closely based on extraction platform instead of sample identity.

**Figure 2.**
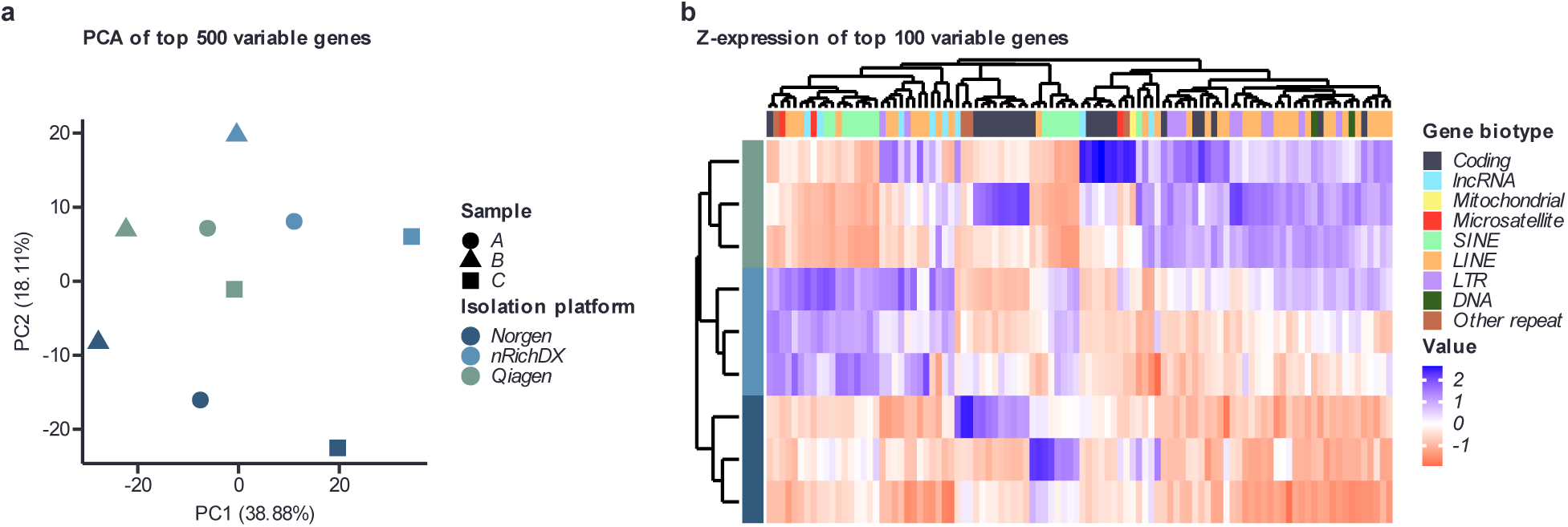
Global cfRNA transcriptome differences between extraction platforms. **(a)** Principal component analysis of the top 500 most variable genes/repeats after variance-stabilized transformation. Colors indicate extraction platform, and shapes indicate plasma pool. **(b)** Heatmap of the 100 most variable genes/repeats. Rows represent extraction platform, and columns show gene/repeat biotype.

To characterize the cfRNA gene biotypes showing the greatest variability across libraries, we performed unsupervised hierarchical clustering of the 100 most variable gene biotypes, including repeat elements (Fig. 2b). Similar to the PCA of the 500 most variable genes, hierarchical clustering grouped libraries by cfRNA extraction workflow rather than by sample of origin, indicating that variation within the highest variance genes was more closely associated with extraction method than with sample identity. Among the 100 most variable genes, 60 genes had the highest mean abundance in Qiagen, 33 had the highest abundance in nRichDX, and 7 were highest in Norgen. Protein-coding (19/22 genes), LINE (23/34), and LTR (9/11) elements were preferentially enriched in Qiagen samples, while SINE (12/15), lncRNA (5/8), and microsatellite (2/3) repeats were most abundant in nRichDX samples.

### cfRNA biotype composition and differential abundances between extraction platforms

To assess whether extraction workflows affected the biotype composition of our cfRNA libraries, reads in each library were grouped by biotype and represented as a percentage of total mapped reads assigned to each biotype (Fig. 3a). To determine the consistency of biotype representation across independent samples, we calculated the mean within-method coefficient of variation in per biotype read percentages for nRichDX (8.6%), Norgen (10.8%), and Qiagen (20.0%).

**Figure 3.**
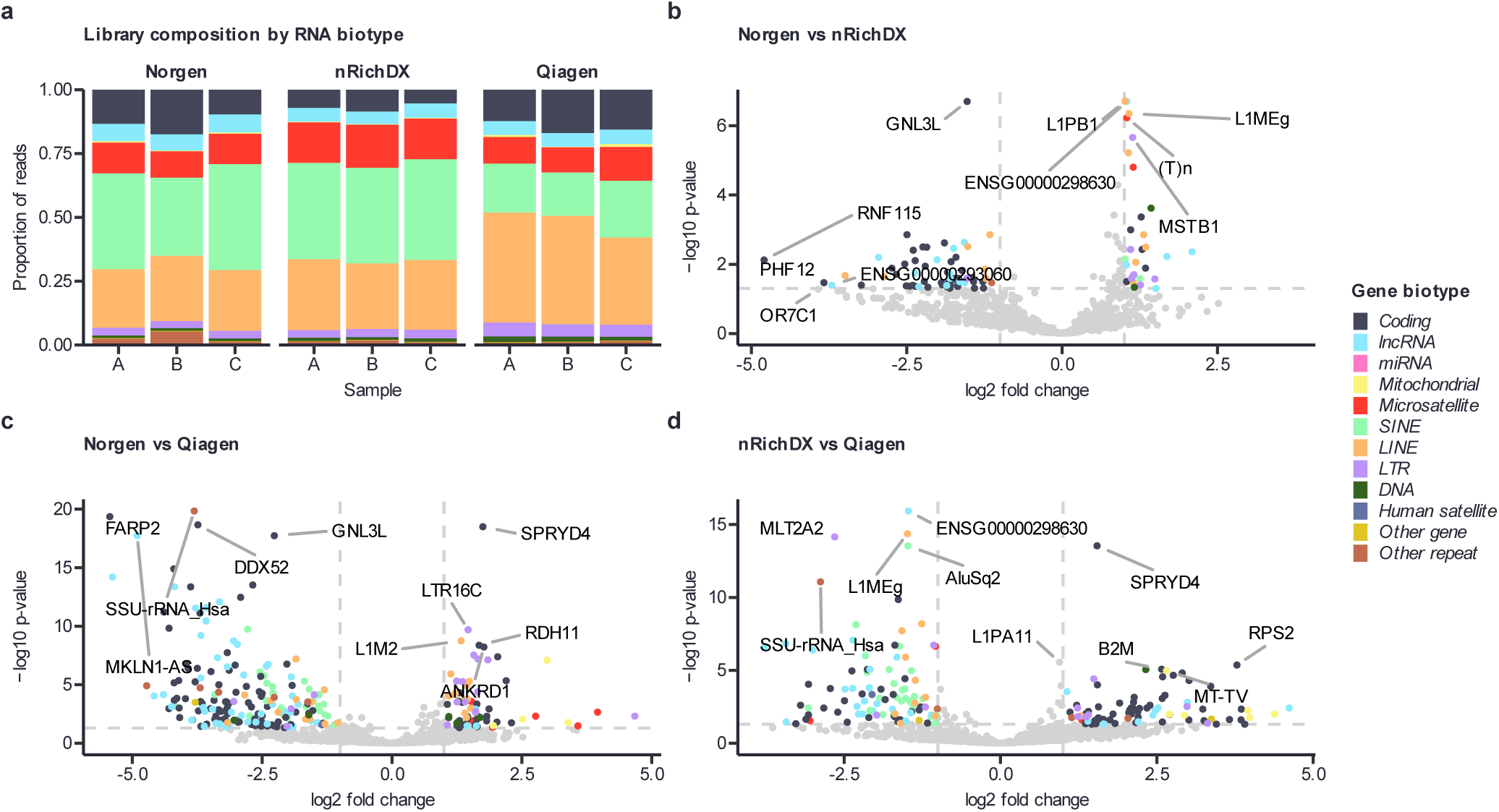
cfRNA biotype composition and differential abundances across platforms. **(a)** Proportion of reads assigned to each RNA biotype for each platform and plasma pool. **(b-d)** Differential expression between extraction platforms. **(a)** Norgen vs. nRichDX, **(b)** Norgen vs. Qiagen, **(c)** nRichDX vs. Qiagen. Each point is a gene or repeat element. The x-axis shows log2fold change, with positive values representing the second listed platform and negative values representing the first listed platform. The y-axis shows the -log10 adjusted p-value. Dashed lines mark |log2fold change| = 1 and adjusted p-value = 0.05. genes/repeats passing both thresholds are colored by biotype, and all others are gray. The top 10 genes/repeats by significance are labeled. Differential expression was determined using DESeq2.

We then performed pairwise differential expression analysis between the cfRNA extraction platforms (Fig. 3b–d). nRichDX versus (vs) Norgen exhibited the fewest significantly differentially expressed genes/repeats (n=92) of all the pairwise comparisons (Fig. 3b). Norgen-extracted cfRNAs were enriched for protein-coding (n=42) and lncRNA (n=12) genes, whereas nRichDX-extracted cfRNAs were enriched for LINE (n=7) and LTR (n=7) repeat element subfamilies. Norgen vs Qiagen had the largest number of significantly differentially expressed genes/repeats (n=253) (Fig. 3c), with Norgen enriched for protein-coding (n=83) and lncRNA (n=45) genes, as well as SINE (n=27) repeat element subfamilies and Qiagen enriched for LTR (n=13) and LINE (n=12) subfamilies, along with a few DNA transposons, microsatellite repeats, and mitochondrial genes. nRichDX vs Qiagen also showed a large number of significantly differentially expressed genes/repeats (n=181), with nRichDX enriched for lncRNA (n=21) genes and SINE (n=28) and LINE (n=10) subfamilies and Qiagen enriched for protein-coding (n=45) genes, together with a few LTR subfamilies and mitochondrial genes.

### Preferential enrichment of gene and repeat biotypes across cfRNA extraction platforms

We separated the pairwise differential expression results by gene/repeat biotype to examine their enrichment differences between cfRNA extraction platforms (Fig. 4). Each platform exhibited preferential enrichment for specific protein-coding (Fig. 4a) and lncRNA (Fig. 4b) genes, including potential biomarker genes with known disease associations. For example, Norgen-extracted cfRNA was enriched for the tumor necrosis factor receptor superfamily member 13C (TNFRSF13C) protein-coding gene [36] relative to both nRichDX (log2-fold change: 2.18, padj = 3.2 × 10⁻³) and Qiagen (log2-fold change: 3.88, padj = 4.4 × 10⁻¹⁴) (Fig. 4a), as well as the p53 upregulated regulator of p53 levels (PURPL) lncRNA gene [37] relative to nRichDX (log2-fold change: 1.57, padj = 0.034) and Qiagen (log2-fold change: 1.92, padj = 9.2 × 10⁻⁵) (Fig. 4b). nRichDX-extracted cfRNA was enriched for the cell-surface transmembrane signaling receptor CD47 protein-coding gene [38] relative to Norgen (log2-fold change: 1.10, padj = 1.0 × 10⁻³) and Qiagen (log2-fold change: 1.63, padj = 1.08 × 10⁻^12^) (Fig. 4a), along with the ADAM metallopeptidase with thrombospondin type 1 motif 9 antisense RNA 2 (ADAMTS9-AS2) lncRNA gene [39] relative to Norgen (log2-fold change: 2.09, padj = 4.4 × 10⁻³) and Qiagen (log2-fold change: 2.10, padj = 1.6 × 10⁻⁴) (Fig. 4b). Qiagen-extracted cfRNA was enriched for the SPRY domain-containing protein 4 (SPRYD4) protein-coding gene [40] relative to Norgen (log2-fold change: 1.75, padj = 3.2 × 10⁻¹⁹) and nRichDX (log2-fold change: 1.54, padj = 2.9 × 10⁻¹⁴) (Fig. 4a).

**Figure 4.**
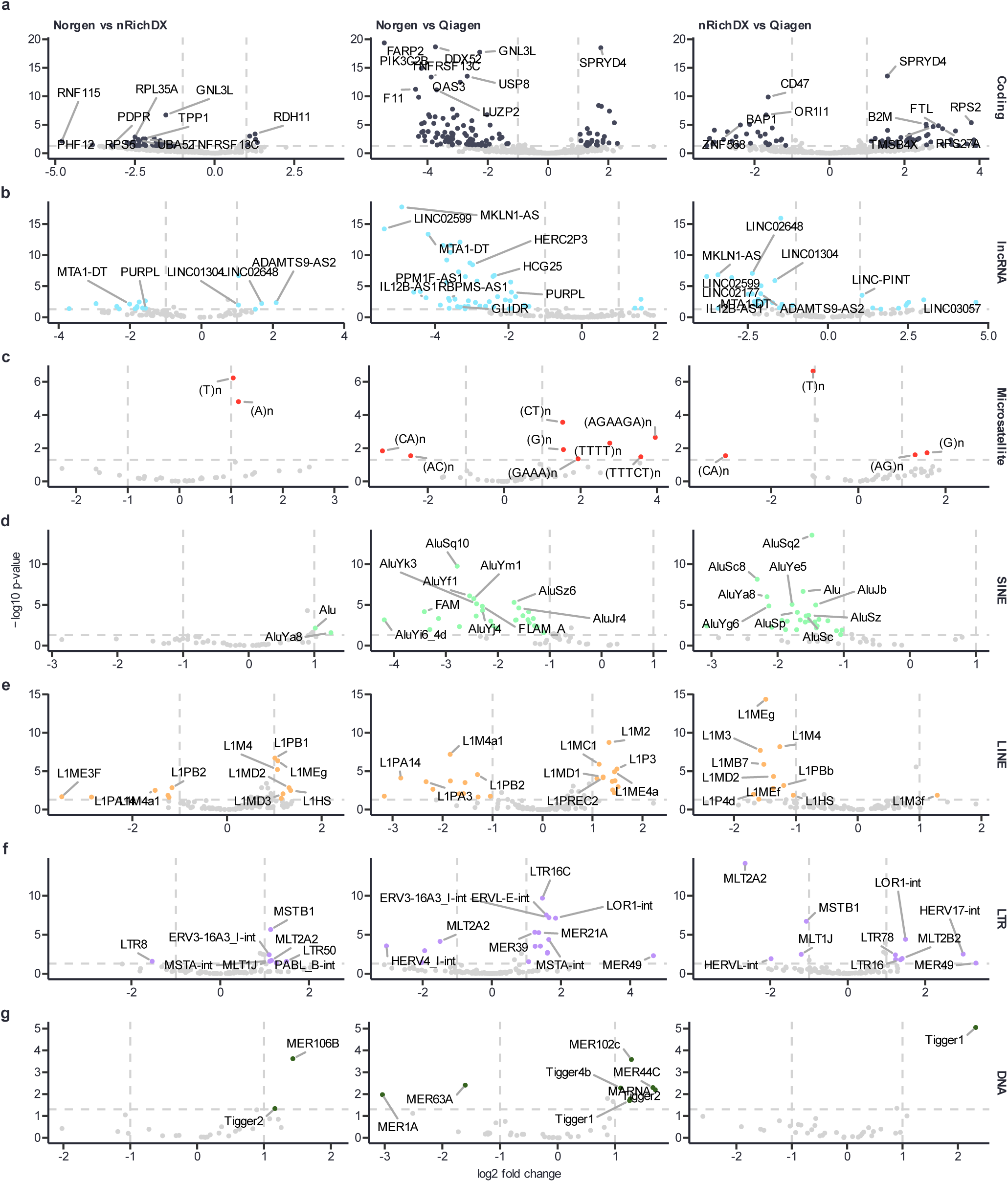
Biotype-specific differences between cfRNA extraction platforms. The differential expression comparisons from Fig. 2a–c, split by biotype. Rows show **(a)** protein-coding genes, **(b)** lncRNAs, **(c)** microsatellites, **(d)** SINEs, **(e)** LINEs, **(f)** LTR retrotransposons and **(g)** DNA transposons. Columns show the three pairwise comparisons: Norgen vs. nRichDX (left), Norgen vs. Qiagen (middle) and nRichDX vs. Qiagen (right). Each point is a gene or repeat subfamily. The x-axis shows log₂ fold change, with positive values indicating higher abundance in the second listed platform. The y-axis shows −log₁₀ adjusted p-value. Dashed lines mark |log₂ fold change| = 1 and adjusted p = 0.05. Features passing both thresholds are colored by biotype (colors as in Fig. 2), and all others are gray. The top ten known genes and repeats by significance are labeled in each panel. (n = 3 pools per platform).

Repeat element subfamilies also showed consistent patterns of enrichment based of cfRNA extraction platform. Microsatellite repeats were preferentially enriched in Qiagen relative to Norgen, with a few repeats also enriched in nRichDX when compared to Norgen (Fig. 4c). SINE subfamilies were significantly enriched in Norgen (n=27 SINE subfamilies) and nRichDX (n=28 SINE subfamilies) relative to Qiagen (Fig. 4d). Moreover, LINE subfamilies were significantly enriched in nRichDX (e.g., L1HS, L1M4, L1MD2, and L1Meg) relative to Qiagen (Fig. 4e).

LTR subfamilies were preferentially enriched in Qiagen (e.g., LOR1-int, LTR78, MER49, and MLT2B2) when compared to Norgen (Fig. 4f), and DNA transposons were also more abundant in Qiagen (e.g., MER102c, MER44C, Tigger4b, and Tigger1) relative to Norgen (Fig. 4g).

## DISCUSSION

While published cfRNA extraction workflows have previously shown preferential recovery of certain biotypes [19,29,31,41,42], these studies focused on well-annotated protein-coding genes or small RNAs, such as microRNAs, and characterized these cfRNAs using short-read sequencing platforms. Here we use long-read nanopore sequencing and repeat-aware analysis on identical plasma samples across three different cfRNA extraction platforms to evaluate technical reproducibility and variability, and to determine the suitability of each platform for extracting a broad range of cfRNA biotypes with disease-relevant biomarker potential.

Although similar amounts of total cfRNA were recovered across the three extraction platforms, each platform exhibited differences in cfRNA read-length distribution, cfRNA biotype composition, and transcript recovery across independent plasma pools. These differences reveal that total cfRNA yield is not indicative of cfRNA extraction quality, as evidenced by the variability we observe in cfRNA transcriptome complexity and reproducibility.

Some of the differences we observe across cfRNA extraction platforms are likely due to the EV enrichment step in the Qiagen platform, whereas the Norgen and nRichDX platforms directly isolate total cfRNA. Given the role of EVs in protecting encapsulated cfRNAs from degradation in the circulation, this may explain the slightly longer cfRNA transcripts and higher N50 values in the Qiagen platform-extracted cfRNAs. However, the cfRNA read lengths were relatively similar in Qiagen vs. nRichDX (cfRNA reads longer than 200 nt: 80% for Qiagen vs. 77.5% for nRichDX) (N50 values: 626 nt for Qiagen vs. 550 nt for nRichDX), suggesting that most of the cfRNAs that are extracted using the nRichDX platform are also likely protected from degradation in EVs. Shorter cfRNAs may also be protected from degradation in ribonucleoprotein complexes [43] and preferentially enriched in Norgen samples, which may explain the lower N50 values in Norgen-extracted cfRNAs (N50: 396 nt).

Beyond broad differences in biotype representation, cfRNA extraction method influenced the measured abundance of individual protein-coding and lncRNA [44] genes, with implications for biomarker detection sensitivity. As representative examples, potential cfRNA biomarkers included *TNFRSF13C*, which encodes the BAFF receptor implicated in antibody deficiency, systemic lupus erythematosus, and B-cell lymphoma [36], *PURPL*, a lncRNA associated with colorectal cancer, liver cancer, and melanoma [37], *ADAMTS9-AS2*, a reported tumor-suppressor lncRNA in gastric, ovarian, bladder, and lung cancers [39], and *SPRYD4*, a reported tumor suppressor in hepatocellular carcinoma and cholangiocarcinoma [40]. Although these disease associations do not establish their utility as circulating biomarkers, their platform-dependent levels in plasma reveal that cfRNA extraction method affects measurement robustness and reproducibility for potential biomarker candidates.

For repeat element-derived cfRNAs, the higher representation of LINE- and LTR-derived cfRNAs in Qiagen samples may reflect the preferential capture of certain EV types by the Qiagen columns, whereas the enrichment of SINE-derived cfRNAs in nRichDX samples may reflect the capture of cfRNAs that are packaged in additional EV classes not captured by Qiagen.

Finally, a notable distinction between cfRNA extraction platforms was their consistency in capturing the same transcripts across plasma pools. nRichDX samples exhibited consistent reproducibility of biotype composition across samples, as well as the largest overlap in the number of consistently detected genes/repeats, whereas Qiagen samples exhibited greater variability across these metrics. This may reflect technical variability but also potential biological variability in the cfRNA contents of secreted EVs. However, one potential source of technical variability is whether a cfRNA extraction workflow is automated (nRichDX) or manual (Qiagen, Norgen). While Qiagen-extracted cfRNA samples had the highest overall correlation across plasma pools, this was driven largely by low-abundance genes/repeats. nRichDX-extracted cfRNA samples exhibited similar correlations when compared to Qiagen sample correlations, but across genes/repeats with consistently higher overall counts, indicating that correlation coefficients may not fully reflect performance differences in consistency and reproducibility between cfRNA extraction platforms.

Taken together, our study provides a benchmark for assessing how extraction platforms shape cfRNA transcriptome representation, as determined by long-read nanopore sequencing, revealing that platform choice is a key determinant of the cfRNA transcripts and biotypes that are reliably and reproducibly captured for biomarker discovery and other clinically relevant RNA liquid biopsy applications.

## METHODS

### cfRNA extraction

Plasma source, processing and study design. Eleven aliquots of human plasma were pooled to create three separate 4 ml plasma pools, which served as biological replicates. Each pool was filtered through a 0.8 µm filter and 1.5 ml of filtered plasma was collected from each replicate. Each filtered replicate was then divided into aliquots for RNA extraction using different protocols on matched input material. Three extraction platforms were evaluated in triplicate: (i) nRichDX (nRichDX Revolution cfTNA Max 20 Kit, Cat. 100499); (ii) Norgen (Plasma/Serum Circulating and Exosomal RNA Purification 96-well kit (Slurry Format), Cat.29500); and (iii) Qiagen (exoRNeasy Maxi Kit, Cat. 77064). After RNA extraction, Zymo Reasearch’s RNA Clean & Concentrator-5 purification (Cat. R1014) and on column DNase I treatment were applied to each sample, according to the manufacturer’s instructions. Library preparation, sequencing, alignment and quantification. For each sample, 2 ng of extracted RNA was reverse-transcribed and barcoded with the Takara Bio SMART-Seq mRNA long-read kit (Cat. 634376), and 80 ng of the resulting cDNA was taken into the Nanopore SQK-LSK114 ligation library preparation. Libraries were sequenced on two Oxford Nanopore PromethION runs using R10.4.1 flow cells (72 h per run): nRichDX and Norgen libraries were pooled on a single flow cell, and Qiagen libraries were run on a separate flow cell.

### Alignment and analysis

Basecalling was performed with Dorado in high accuracy mode using the model. Reads were aligned to the GRCh38 primary assembly using a RepeatMasker annotation with minimap2 (v2.26) [45] using the -ax splice:sr preset. Mapped reads were then quantified using Oarfish [46]. A minimum-expression filter of rowSums(count_matrix >= 3) >= 3 was applied to the gene-level matrix. Bioinformatic analysis was performed in R (v4.4.3). Quantification output from oarfish was imported into R using tximeta (v1.24.0) [47], and library normalization and differential expression analysis was performed using DESeq2 (v1.46.0) [48]. Genes with less than three reads in more than 6 samples were filtered out prior to analysis. Read lengths from the generated libraries were calculated using the aligned reads after adapter removal and low-quality base trimming. Spearman correlation coefficients were calculated using the log10 normalized quantifications for each gene to lower the rate of heteroskedasticity. Differential expression fold-changes were reduced using the apeglm [49] method, and the p-values for each gene were corrected using Bonferroni correction. Significance was defined as any gene with a reduced log2FoldChange greater than 1, and an adjusted p-value lower than 0.05. Principal component analysis was performed using the normalized quantifications of the top 500 genes with the greatest variance. The heatmap of the samples was calculated using the Z-score of the normalized quantifications, with the Z-score calculated at a per-gene level. Hierarchical clustering was performed using the Z-scores of the top 100 most variables genes, and the branches were ordered by baseMean expression of the genes in each cluster.

## DATA AVAILABILITY

Sequencing data generated in this study have been deposited in NCBI GEO.

## ACKNOWLEDGEMENTS

We thank current and former members of the Kim Lab for helpful discussions and support, including Piper Mahn and Delaney Boman for their assistance.

## AUTHOR CONTRIBUTIONS

D.H.K. conceived and designed the study, C.S.M. performed all experiments, C.S.M. and A.D.H. performed computational analyses and generated figures, D.H.K. supervised research, and C.S.M and D.H.K. wrote the paper with input from A.D.H.

## ETHICS DECLARATIONS

### Ethics Statement

De-identified human plasma samples that were collected with informed consent were purchased from BioIVT.

## REFERENCES

1. Anfossi, S., Babayan, A., Pantel, K. & Calin, G. A. Clinical utility of circulating non-coding RNAs - an update. Nat Rev Clin Oncol 15, 541–563 (2018).

2. Lo, Y. M. D. & Chiu, R. W. K. The Biology and Diagnostic Applications of Plasma RNA. Annals of the New York Academy of Sciences 1022, 135–139 (2004).

3. Koh, W. et al. Noninvasive in vivo monitoring of tissue-specific global gene expression in humans. Proc Natl Acad Sci U S A 111, 7361–6 (2014).

4. Larson, M. H. et al. A comprehensive characterization of the cell-free transcriptome reveals tissue- and subtype-specific biomarkers for cancer detection. Nat Commun 12, 2357 (2021).

5. Reggiardo, R. E. et al. Profiling of repetitive RNA sequences in the blood plasma of patients with cancer. *Nat*. Biomed. Eng 7, 1627–1635 (2023).

6. Loy, C., Ahmann, L., De Vlaminck, I. & Gu, W. Liquid Biopsy Based on Cell-Free DNA and RNA. Annual Review of Biomedical Engineering 26, 169–195 (2024).

7. Toden, S. & Goel, A. Non-coding RNAs as liquid biopsy biomarkers in cancer. Br J Cancer 126, 351–360 (2022).

8. Schwarzenbach, H. & Pantel, K. Circulating DNA as biomarker in breast cancer. Breast Cancer Res 17, 136 (2015).

9. Delgado, P. O. et al. Characterization of cell-free circulating DNA in plasma in patients with prostate cancer. Tumour Biol 34, 983–986 (2013).

10. Reggiardo, R. E. et al. Epigenomic reprogramming of repetitive noncoding RNAs and IFN-stimulated genes by mutant KRAS. bioRxiv 10.1101/2020.11.04.367771 (2020) doi:10.1101/2020.11.04.367771.

11. Reggiardo, R. E. et al. Mutant KRAS regulates transposable element RNA and innate immunity via KRAB zinc-finger genes. Cell Reports 40, 111104 (2022).

12. Tsang, J. C. H. et al. Integrative single-cell and cell-free plasma RNA transcriptomics elucidates placental cellular dynamics. Proc Natl Acad Sci U S A 114, E7786–E7795 (2017).

13. Vorperian, S. K. et al. Cell types of origin of the cell-free transcriptome. Nat Biotechnol 40, 855–861 (2022).

14. Welsh, J. A., et al. Minimal information for studies of extracellular vesicles (MISEV2023): From basic to advanced approaches. J of Extracellular Vesicle 13, e12404 (2024).

15. Wang, J., Ma, P., Kim, D. H., Liu, B.-F. & Demirci, U. Towards microfluidic-based exosome isolation and detection for tumor therapy. Nano Today 37, 101066 (2021).

16. Dixson, A. C., Dawson, T. R., Di Vizio, D. & Weaver, A. M. Context-specific regulation of extracellular vesicle biogenesis and cargo selection. Nat Rev Mol Cell Biol 24, 454–476 (2023).

17. Valadi, H. et al. Exosome-mediated transfer of mRNAs and microRNAs is a novel mechanism of genetic exchange between cells. Nat Cell Biol 9, 654–659 (2007).

18. Kim, H. J. et al. Selective enrichment of plasma cell-free messenger RNA in cancer-associated extracellular vesicles. Commun Biol 6, 885 (2023).

19. Srinivasan, S. et al. Small RNA Sequencing across Diverse Biofluids Identifies Optimal Methods for exRNA Isolation. Cell 177, 446–462.e16 (2019).

20. Moufarrej, M. N. et al. Early prediction of preeclampsia in pregnancy with cell-free RNA. Nature 602, 689–694 (2022).

21. Ngo, T. T. M. et al. Noninvasive blood tests for fetal development predict gestational age and preterm delivery. Science 360, 1133–1136 (2018).

22. Toden, S. et al. Noninvasive characterization of Alzheimer’s disease by circulating, cell-free messenger RNA next-generation sequencing. Sci. Adv. 6, eabb1654 (2020).

23. Wang, H. et al. Depletion-assisted multiplexed cell-free RNA sequencing reveals distinct human and microbial signatures in plasma versus extracellular vesicles. Clinical & Translational Med 14, e1760 (2024).

24. Wijeratne, S. et al. Full-length isoform concatenation sequencing to resolve cancer transcriptome complexity. BMC Genomics 25, 122 (2024).

25. Tan, X. et al. Long-Read Sequencing Reveals RNA Splicing Complexity in Human Diseases. Comput Struct Biotechnol J 35, 0052 (2026).

26. Katopodi, X.-L., Begik, O. & Novoa, E. M. Toward the use of nanopore RNA sequencing technologies in the clinic: challenges and opportunities. Nucleic Acids Research 53, gkaf128 (2025).

27. Peddu, V. et al. RNA liquid biopsy via nanopore sequencing for novel biomarker discovery and cancer early detection. bioRxiv 2025.07.02.662774 (2025) doi:10.1101/2025.07.02.662774.

28. Lau, B. T. et al. Single-molecule methylation profiles of cell-free DNA in cancer with nanopore sequencing. Genome Med 15, 33 (2023).

29. Tuñí-Domínguez, C. et al. Systematic cross-study assessment of RNA-Seq experimental workflows for plasma cell-free transcriptome profiling. 2025.07.10.664092 Preprint at 10.1101/2025.07.10.664092 (2025).

30. Bao, P. et al. Peak analysis of cell-free RNA finds recurrently protected narrow regions with clinical potential. Genome Biol 26, 119 (2025).

31. exRNAQC Consortium. Blood collection tube and RNA purification method recommendations for extracellular RNA transcriptome profiling. Nat Commun 16, 4513 (2025).

32. Makarova, J., Maltseva, D. & Tonevitsky, A. Challenges in characterization of transcriptomes of extracellular vesicles and non-vesicular extracellular RNA carriers. Front. Mol. Biosci. 10, 1327985 (2023).

33. Carrillo, D. et al. Transposable element RNA dysregulation in mutant KRAS(G12C) 3D lung cancer spheroids. bioRxiv 10.1101/2023.02.27.530369 (2023).

34. Khojah, R. et al. Extracellular RNA signatures of mutant KRAS(G12C) lung adenocarcinoma cells. 2022.02.23.481574 bioRxiv 10.1101/2022.02.23.481574 (2022).

35. Tian, B. A large-scale analysis of mRNA polyadenylation of human and mouse genes. Nucleic Acids Research 33, 201–212 (2005).

36. Budde, L. E. et al. BAFF-R CAR T-cell therapy for relapsed or refractory B-cell lymphomas. Lancet 408, 891–893 (2026).

37. Li, X. L. et al. Long Noncoding RNA PURPL Suppresses Basal p53 Levels and Promotes Tumorigenicity in Colorectal Cancer. Cell Rep 20, 2408–2423 (2017).

38. Majeti, R. et al. CD47 Is an Adverse Prognostic Factor and Therapeutic Antibody Target on Human Acute Myeloid Leukemia Stem Cells. Cell 138, 286–299 (2009).

39. Liu, W., Luo, W., Zhou, P., Cheng, Y. & Qian, L. Bioinformatics Analysis and Functional Verification of ADAMTS9-AS1/AS2 in Lung Adenocarcinoma. Front Oncol 11, 681777 (2021).

40. Ma, Z. et al. Identification of SPRYD4 as a tumour suppressor predicts prognosis and correlates with immune infiltration in cholangiocarcinoma. BMC Cancer 23, 404 (2023).

41. El-Khoury, V., Pierson, S., Kaoma, T., Bernardin, F. & Berchem, G. Assessing cellular and circulating miRNA recovery: the impact of the RNA isolation method and the quantity of input material. Sci Rep 6, 19529 (2016).

42. Cabús, L., Lagarde, J., Curado, J., Lizano, E. & Pérez-Boza, J. Current challenges and best practices for cell-free long RNA biomarker discovery. Biomark Res 10, 62 (2022).

43. Arroyo, J. D., et al. Argonaute2 complexes carry a population of circulating microRNAs independent of vesicles in human plasma. Proc. Natl. Acad. Sci. U.S.A. 108, 5003–5008 (2011).

44. Reggiardo, R. E., Maroli, S. V. & Kim, D. H. LncRNA Biomarkers of Inflammation and Cancer. in Long Noncoding RNA (ed. Carpenter, S.) vol. 1363 121–145 (Springer International Publishing, Cham, 2022).

45. Li, H. Minimap2: pairwise alignment for nucleotide sequences. Bioinformatics 34, 3094–3100 (2018).

46. Zare Jousheghani, Z., Singh, N. P. & Patro, R. Oarfish : enhanced probabilistic modeling leads to improved accuracy in long read transcriptome quantification. Bioinformatics 41, i304–i313 (2025).

47. Love, M. I. et al. Tximeta: Reference sequence checksums for provenance identification in RNA-seq. PLoS Comput Biol 16, e1007664 (2020).

48. Love, M. I., Huber, W. & Anders, S. Moderated estimation of fold change and dispersion for RNA-seq data with DESeq2. Genome Biol 15, 550 (2014).

49. Zhu, A., Ibrahim, J. G. & Love, M. I. Heavy-tailed prior distributions for sequence count data: removing the noise and preserving large differences. Bioinformatics 35, 2084–2092 (2019).

